# A validation approach for computational models of TMS induced brain currents using motor evoked potentials

**DOI:** 10.1101/2023.02.24.528183

**Authors:** Petar I. Petrov, Jord Vink, Stefano Mandija, Nico A.T. van den Berg, Rick M. Dijkhuizen, Sebastiaan F.W. Neggers

**Affiliations:** Dept of Psychiatry, Brain Center Rudolf Magnus, University Medical Center Utrecht, 3508 GA, Utrecht, the Netherlands; Biomedical Magnetic Resonance Imaging and Spectroscopy Group, Center for Image Sciences, University Medical Center Utrecht and Utrecht University, Utrecht, The Netherlands; Dept of Radiotherapy, University Medical Center Utrecht, 3508 GA, Utrecht, the Netherlands

## Abstract

The adoption of transcranial magnetic stimulation (TMS) has steadily increased in research as a tool capable to safely and non-invasively stimulate both the central and peripheral nervous systems. Initial clinical applications were limited to diagnostic use of TMS and readout signals such as electromyograms (EMG). Subsequently, repetitive TMS (rTMS) was appreciated for its therapeutics benefits as well. However, even after a decade of use of rTMS as an alternative treatment of major depression disorder in psychiatry, the mechanism of action is still not well understood. Computer models predicting the induced electric field distribution in the brain have been suggested before in the hope to resolve at least some of the uncertainty and resulting variable treatment response associated with the clinical use of TMS.

We constructed a finite element model (FEM) of the head using individual volumetric tissue meshes obtained from an MRI scan and a detailed model of a TMS coil that together can predict the current induced in the head of a patient at any given location with any given coil position and orientation. We further designed several potential metrics of how a TMS induced current induced neuronal activation in the motor cortex, and added this to the model. We validated this model with motor evoked potentials (MEPs), EMG responses of the hand muscles after TMS on the motor cortex, in an experiment on 9 healthy subjects. We adopted a tailored MEP mapping protocol for model validation, which unlike traditional grid mappings, varies the TMS machine output intensity between stimulation locations. We further varied coil orientation on each point stimulated to allow exploration of the angular dependency of the model MEPs. Taken together, this approach covers a wide domain and scope of the modeled and measured responses, which are optimally suited for model validation. For each subject the motor hotspot was carefully identified using individual cortical anatomy and BOLD fMRI measurements.

Modeled activation in the motor cortex did not show a good correlation to the observed magnitude of the observed MEPs, for none of the neuronal activation metrics adopted. For an activation metric that was asymmetric, taking into account induced current direction with respect to the motor cortex sulcal wall, was marginally better than other metrics. Generally all activation metrics based on induced currents performed better than a control metric agnostic of induced electric field magnitude. Our results suggest that one should take into account components of the injected currents and their relationship to the morphology of the underlying motor cortex, but the coarse metrics we used to model the relationship between induced current and neuronal activation probably did not do justice to the complex neuronal circuitry of the cortical sheet. Furthermore, it seemed MEP magnitudes in our experiment are too variable over subsequent stimulations, which could be mitigated by more repetitions per stimulation location and orientation.

Further efforts to construct validated models predicting TMS effects in individual patients brains should incorporate microcircuits interactions in the cortical sheet, in addition to induced electrical field models, and take into account inherent trial to trial variability of MEPs.

## 1 Introduction

TMS is a non-invasive brain stimulation (NIBS) and modulation technique. Based on the physical principle of electromagnetic induction, it generates magnetic fields, that freely penetrate the skull and can inject relatively focal cortical currents with the appropriate coil design. TMS is increasingly used to treat diseases of the central nervous system, obtain diagnosis after brain trauma, and investigate the organization of the brain [1].

However, how exactly these currents flow and are shaped by non-isotropically conducting brain morphology, and subsequently interact with or modulate neuronal activation, is still an open question. Computer based models predicting TMS-induced currents and subsequent elicited activation have been an active topic of research for more than two decades from which significant progress in understanding TMS effects has been made.

Due to recent advances in computational science, it is now viable, affordable, and possible to model the interaction of electric and magnetic fields with human brain tissue in impressive detail. Initial models of the human cortex were limited to highly idealized geometrical shapes representing the different conductive tissues in the head, and the application of finite element modeling of the incident and induced fields to such compartments. For example, spherical models with superficially derived sub-layers for each major tissue with realistic radius values to account for each tissue thickness have been attempted in the past [2,3]. Later, more sophisticated models were developed where the highly inhomogeneous and anisotropic properties typical for the human brain are captured in sufficient detail, thanks in part to advanced MR imaging. Such models often consist of several separate compartments, segmented automatically or semi-automatically, from T1-weighted anatomical scans generating maps of major brain tissues such as gray matter (GM), white matter (WM), cerebral spinal fluid (CSF), skull, and skin. To approximate the conductive properties of the highly anisotropic white matter, several groups have successfully utilized diffusion-weighted imaging to introduce anisotropic conductivity per mesh element making such models even more realistic [4].

However, despite such progress and the further sophistication of computational modeling of inductively induced currents, only a very small number of such models have been empirically validated. At present, it is unclear what assumptions and simplifications that are unavoidable for any computational model of conductive properties of live tissue, are valid, and which are not. Hence it is not clear to what extent predicted neuronal currents can be relied upon for clinical use. For the incident field or the portion of the electric field induced by the mere geometry of the coil, a few successful validations have been published that confirmed such models at least partially, including our group. By using elaborate MRI-compatible TMS setups, phase accumulation models based on modeled incident TMS evoked currents were validated quite successfully with deviations of only 1– 5% in phase accumulation maps (proportional to incident current magnitude), thus validating this portion of the model for total E-field induced in the head successfully [5]. The situation is much less fortunate for the largest part of such individual TMS-induced activation models. The second part of the total E-field, governed by the inhomogeneity of tissues in the head causing current accumulation and requiring FEM to model, seldom gets empiricaly validated. A notable exception has been the work of Bungert et a [6–8] where EMG and fMRI were used to validate the responses in the muscle or directly in the brain, respectively. Although some correspondence with measured responses was established, a large portion of unexplained variance remained. Only a small portion of TMS modeling studies have looked into validating their results empirically.

Whereas Bungert et al used varying coil orientations to obtain modeled and measured responses in a varying domain, in their protocols the coil was rotated in small steps while being fixated on one location, and only shifted laterally. Thus missing possible more anterior or posterior coil positions that limits the scope and variability of the modeled versus measured responses. Alternatively, although not used in FEM validations to our knowledge, more classic grid mapping studies using a large number of stimulation locations a rectangular grid in combination with one or a few discreet coil orientations have been published to explore response maps of the motor cortex [9–11]. These studies have provided insights into the spatial distribution of TMS-evoked MEPs but were not used for current model validation. A disadvantage of such experimental designs could be overly long sessions due to the number of TMS discharges required, which would result in MEP amplitude habituation (a.k.a. repetition suppression) [12], making it less suitable for current model validation.

In this study, we developed multi tissues 3D finite element model (FEM) of TMS-evoked currents in the brain, based on a detailed coil model and derived from subject-specific anatomical MR images. To predict the MEP amplitudes at several stimulation locations, we propose several hypotheses (activation ‘metrics’) on how such currents can evoke neuronal activation in the cortical sheet. These metrics include a simple amplitude of the induced current integrated over a certain volume of tissue and several more sophisticated metrics with angular dependencies of local activation on induced current direction (see paragraphs below and the methods section for details). The metrics are motivated by the architecture of the microcircuitry in the cortical sheet. We also tested two ideas on where in the pre-central gyrus MEPs measured at the thumb are primarily evoked: only the posterior wall or the entire gyrus including the premotor cortex. To validate our modeling approach, we conducted experiments with mono-phasic TMS in single pulse mode on the motor cortex (hand area) in combination with EMG on the thumb and index finger abductor muscle (first dorsal interosseous) of each subject.

Importantly, we propose a novel more optimal, and comprehensive approach to probe and map the MEP response from the motor cortex for TMS current modeling validation. In this study we adopted stimulation locations evoking MEPs in a cross-shaped layout, covering Central, Posterior, Anterior, Medial and Lateral locations relative to the hand knob as defined by individual functional fMRI maps of finger movements [13], while probing coil orientation in the four major orthogonal directions. We chose the orientations such that we could investigate possible current direction dependent as well as position effects, covering an optimal domain of spatial parameters.

Many FEM TMS modeling studies produced predicted currents in the shape of 3D maps [14,15], which is not the same as neuronal activation. As discussed above, when validating such models with empirical measures such as TMS-evoked fMRI or MEPs, one needs a metric that reflects neuronal activation as a result of local current estimates, to bridge the gap between the neuronal or muscle responses to TMS and the evoked currents in the brain. Commonly, the maximum electrical field magnitude at each location in a stimulated area is reported in the literature as a proxy for ‘neuronal activation’ [16]. Others have adopted an activation metric proportional to the direction of the electric field relative to the folding cortical surface. Most notably the C3 metric proposed by Fox et al [17], based on a cortical column model, where orthogonal orientation to the outer GM surface fields is suggested as most effective and as such most pronounced to the physiological effect of stimulation. The same model was demonstrated numerically (FEM) and supported with functional (H_2_^15^O) PET imaging, as evidence for this hypothesis [18]. Recently, however, such a simplistic generalization of neuronal responses to local current patterns was challenged by the observation that the site of activation is not where we would expect it. Based on FEM models combined with the aforementioned C3 metric [6], albeit through indirect evidence of comparing results from a model to previously reported functional imaging studies, Bungert et al [6] has confronted the notion that any prime component of the E-field relative to the outer cortical layer can be a good estimator alone. Rather they proposed the relative standard deviation (SD) of the product across several coil orientations at a fixed site as a biophysical estimate for the potential MEP response.

We, therefore, did not just focus on this single and challenged metric (C3) but included several potential metrics mimicking the link between local modeled current patterns and compared the results of our FEM EM simulations against all proposed metrics. The metrics we developed take into account the electric field amplitude in combination with its tangential and radial components relative to the targeted cortical surface. Finally, we adopted an additional very simplistic metric not depending on induced current in any way, but only on a distance from the electric field maximum to a stimulation point-target, which can serve as a H0-hypothesis. Including such a zero-hypothesis metric allowed us to test the validity of some naive approaches that are completely independent of the induced E-fields, rather they rely on simple distance from coil to target adjustments to estimate the recommended dosimetry [19]. Previous work by [20], has already challenged such approaches, still, we tried to adequately account for such field-independent predictors in our comparison.

Finally, validation of TMS-evoked current models with MEPs is an indirect approach, even when we manage to establish a good metric for the relationship between induced current models and local neuronal activation. Also, evoked MEPs further rely on assumptions of how the neuronal activation pattern travels from the stimulated motor cortex through the cortico-spinal tract and ultimately innervates the muscles, measured by EMG. We evaluate the numerically produced modeled E-fields on two regions of interest (ROIs) around the hand knob in the motor cortex, one representing the anterior wall of the central suclus, and the other covering the entire precentral gyrus around the hand knob, also including the pre-motor cortex, similar to Bungert et al [6]. With this, we also tested assumptions about what parts of the tissue in the motor cortex, in which currents are evoked, contribute to the magnitude of the signal arriving at the muscle through the corticospinal tract.

## 2 Materials and Methods

### 2.1 Participants

For this and another study, a total of 11 subjects were recruited. Data from 9 subjects (4 males and 5 females) were processed and analyzed. The experimental procedure was approved by the medical ethical committee of the University Medical Center Utrecht (UMCU), Utrecht, The Netherlands (protocol 16-469/D).

All participants were included only under written consent and without any counterindications to TMS reported (personal and/or family history of epilepsy; not currently on any medication). Also, all participants were right-handed. Two of the participants dropped out of the study after failing to follow some of the sessions.

### 2.2 Experimental Setup

#### 2.2.1 MR Image Acquisition

All MR acquisitions were performed with a 3T MR scanner Achieva (Phillips, The Netherlands). Anatomical T1 weighted scan was acquired with a TR/TE of 10.015/4.61ms, a flip angle of 8°, voxel size of 0.75×0.75×0.8mm, scan duration of 677s, 225 slices with a slice gap of 0 mm. For fMRI time series measurements, a single-shot EPI sequence was acquired with 250 dynamics, a TR/TE of 2,000/23ms, a flip angle of 70, a voxel size of 4×4×4 mm, 30 slices with a slice thickness of 3.6mm and a slice gap of 0.4mm.

#### 2.2.2 Electromyography (EMG)

A Neuro-MEP-4 (Neurosoft, Russia) EMG device with 4 channels was used, 20 kHz sampling rate with an amplification gain factor of 1000 (up to 60mV). The software with the device supplies impedance monitoring, which we used to keep impedance low (<10kΩ) in the green zone (green/yellow <25kΩ; yellow/red <40kΩ). We used surface Ag/AgCl electrodes (FIAB, Italy) (REF: F3001ECG), which were placed over the right hand first dorsal interosseous (FDI), abductor digiti minimi (ADM) and extensor carpi radialis (ECR) muscle in a belly-tendon montage and the ground/reference electrode was attached to the wrist of the left hand. For this study, only channel-1 (FDI) traces were processed and analyzed.

#### 2.2.3 Transcranial Magnetic Stimulator (TMS)

The stimulator we used was the Neuro-MS Monophasic TMS (Neurosoft, Russia) with a figure-8 flat coil (Reference: FEC-02-100) in a single pulse mode. All sessions were navigated and EMG MEPs responses were recorded using the Neural Navigator software in the “motor mapping” variant, version 3.1 (Brain Science Tools B.V, The Netherlands). All participants were asked to sit in a relaxed manner on a normal chair with the palm facing upward resting on a table surface, while the head was placed on a chin-support frame supplied by Brain Science Tools BV to constrain head movements.

#### 2.2.4 Computational FEM Electro-magnetic Solution

To compute the electromagnetic effects of TMS, we relied on a detailed Finite Element Model (FEM) of the head, which depends on accurate coil field calculation and the subsequent secondary field estimate. Commonly, an iterative linear optimization numerical routine is used, the gradient descent method [21]. The secondary field is due to charge accumulation in a highly non-homogeneous medium and is super-exposed along with the primary field on the domain, acting in opposite direction and generally reducing the total electric field, according to the formula:

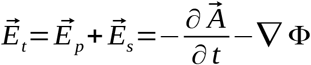, where A is the magnetic vector potential and F is the electrical scalar potential. While the first term on the right-hand side of this formula is entirely governed by the temporal discharge characteristics of the stimulator and the geometry of the coil, forming a classic CLR circuit, the second term is fully dependent on the dielectric properties of the conductive medium that is stimulated. The two fields are often referred to as the magnetic vector potential and scalar gradient potential, respectively. To numerically derive those we developed a custom in-house module to compute the coil fields as an addition to SCIRun which is an integrated virtual environment for interactive FEM solving and visualization. ^1^

#### 2.2.5 Computational Coil Modeling

It is important to have a good approximation of the primary (incident) E-field induced by the coil alone. We expected it to be the primary contributor to the shape and to a great extent the magnitude of the injected cortical currents. In some of our previous work, we investigated the accuracy of the thin-wire solver based on the BiotSavart method used in the current study. We conducted B-field mapping experiments on phantoms inside an MR scanner, utilizing phase accumulation imaging as the results of external magnetic fields inside the scanner [5,22]. Our results pointed to a clear 20% E-field estimate discrepancy of the idealized geometries versus more realistic multi-windings ones, such as single thick wire per coil windings in the case of figure-8 TMS coil, whereas coil models taking into account spiral winding geometry of actual TMS coils only showed discrepancies in the range of 1-5% compared to measurements.

Hence, coil geometry was modeled as a single layer of 10 windings, in the form of circular rings, with R most inner 15mm and R most outer ring 55mm. To generate the geometry we developed a custom SCIRun module, which was previously published [5].

The Neuro-MS mono-phasic stimulator is capable of delivering currents up to 12.52 kA thanks to a powerful capacitor that allows charging up to 2800 V at t=0, considering 100% MO (Machine Output) and the formula 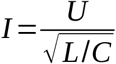. When discharged, it produces a short pulse of 0.2 ms duration and 0.05 ms ramp time for the current to go from 0 to I_max, resulting in max dI/dt of 250×1E^6 [A/s], which we employed to deliver an estimate of the incident electric field dA/dt using the Biot-Savart numeric approximation.

#### 2.2.6 Human Head Modeling

It is critical for the quality of our FEM model to use an anatomically correct model of the human head with additional geometry constraints that guarantees qualitative numeric results. For that reason, we added an elaborate image-to-3D geometry pipeline, from segmentation to final mesh generation.

Semi-automatic tissue segmentation from anatomical T1-weighted MR image was conducted for each participant using SPM12 (FIL, UCL, London) and the unified segmentation method [23]. Five tissue types were classified: CSF, GM, WM as well as skull and scalp. The output of SPM12 is in the form of probability maps one per tissue class, which we further thresholded to produce a single combined binary segmentation file with minimal additional manual corrections needed. As the tissue classification maps produced by SPM for scalp and skull did not have a well-defined border but rather a grainy transition, we decided to merge scalp and skull volumes.

Some manual cleaning around some participants’ M1 area was necessary to eliminate the accidental merging of two adjacent gyri or to compensate for a very thin CSF layer that might introduce holes where the skull directly touches the GM. For a clear example of such manual cleaning around the GM we refer to figure 1-a.

**Figure 1:**
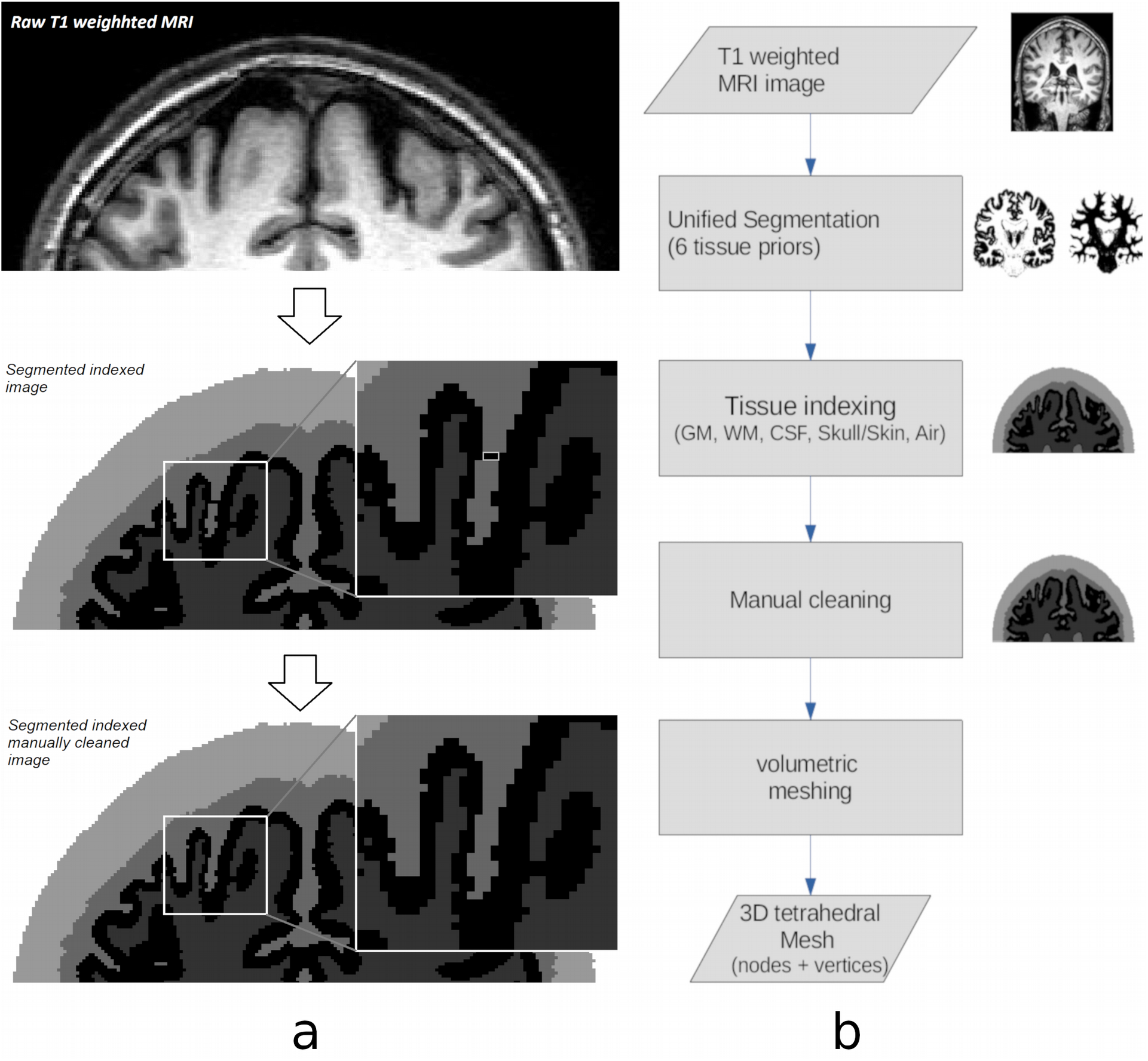
On the left (a): From top to bottom a coronal cut from anatomical T1 weighted MRI to binary partitioned tissue classification map. On the right (b): Typical steps involved in image processing pipelines from raw scanner images to final volumetric 3D model.

In addition to correcting for artifacts, we aimed to produce as smooth as possible surface boundaries for each compartment. Once again we took care not to compromise the final mesh geometry due to a too aggressive smoothing procedure, which invol’ed an iterative routine of alternating inflation-deflation operations in image space, step 1 (per tissue compartment isolation) and 2 (image smoothing) of BioMesh3D ^2^ [24]. Those images were fetched to our meshing algorithm, again one per tissue type to produce the final FEM mesh. Mesh generation was done with Cleaver2 ^3^. The cleaving routine is guaranteed to generate tetrahedral elements within some quality limits that avoid some geometries unfit for numerical evaluation [25–27].

A conceptual diagram of the main steps taken from anatomical T1-weighted images to the final FEM mesh generated is demonstrated as a pipeline with several sequential major processing steps, in figure 1-b. The following conductivity sigmas were used for the FEM: Skull: 0.002[S/m]; White matter: 0.12[S/m]; Gray matter: 0.27[S/m]; CSF: 1.67[S/m] all within the range of previously reported values from the literature [28].

#### 2.2.7 Activation area ROI

To validate our TMS induced current model with TMS evoked MEPs, we have to first answer two questions on how exactly TMS induced currents lead to a MEP. First, we must have assumptions on how currents generate neuronal activations, and, second, we must define how this neuronal activation contribute to a muscle signal through the cortico-spinal tract. In this section, we explain how we construct two possible regions of interest around the hand knob whose summed neuronal activation could contribute to thumb MEP. In the next section, we hypothesize how ‘activation’ could arise in the neuronal circuits contained in the cortical sheet evoked by the TMS-induced currents.

To obtain the two possible regions of interest (ROIs) in which activation can be expected to contribute to an MEP, inside SCIRun we thresholded the resulting fMRI map until we saw a clear representation in the motor area controlling the hand, anatomically referred to as the “hand knob” (see table 1 for thresholded values). Two surface patches were isolated from the pial surface (the outer gray matter-CSF boundary), see figure 2. The extent of each surface was hand-picked to always span the crown of the precentral gyrus. For the larger ROI-1, the anterior wall of the precentral gyrus facing the precentral sulcus and encompassing the so called ‘premotor’ cortex is included as well as the posterior wall of the precentral gyrus, facing the central sulcus. It hence encompasses the entire pre-central gyrus around the hand knob. For the smaller ROI-2, only the posterior wall of the pre-centrul gyrus around the hand knob is included, thus encompassing the classical primary motor cortex. See figure 2 for a visualization of ROI-1 and ROI-2 of one of the participants.

**Table 1:** Participants and Meshing Parameters.

| # | Subject | Age | Sex | RMT |
| --- | --- | --- | --- | --- |
| 1 | SU01 | 21 | F | 41 |
| 2 | SU02 | 34 | M | 40 |
|  | SU03 | 25 | F | 39 |
| 3 | SU04 | 18 | M | 34 |
| 4 | SU05 | 19 | F | 40 |
| 5 | SU06 | 24 | F | 38 |
| 6 | SU07 | 23 | M | 40 |
| 7 | SU08 | 20 | F | 41 |
| 8 | SU09 | 19 | F | 42 |
|  | SU10 | 20 | F | 41 |
| 9 | SU11 | 46 | M | 43 |

**Figure 2:**
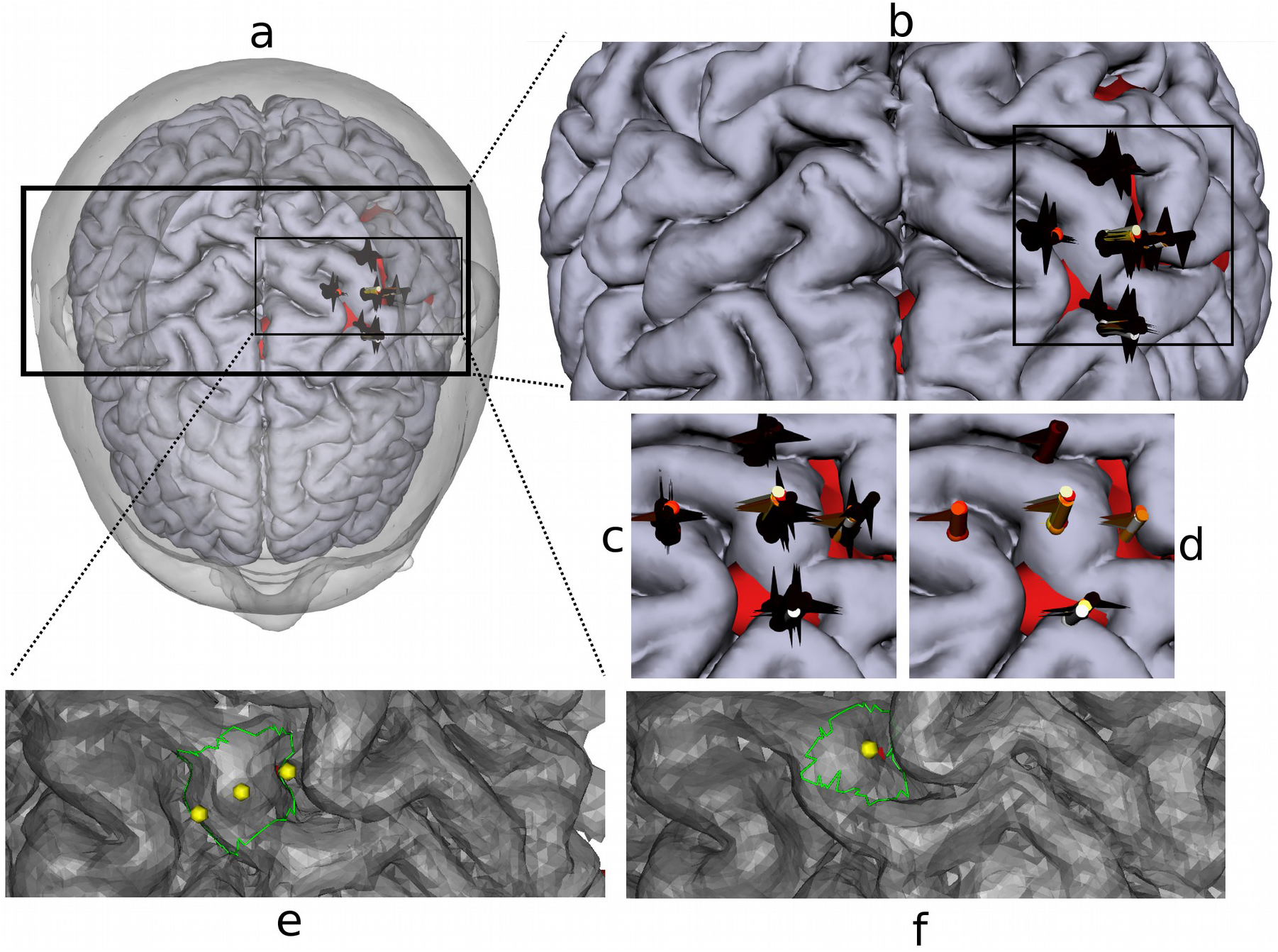
Demonstration of a completed navigation and cross-mapping session with subject SU06 with screen captures from the navigation software we used. (a) an axial from-top view of 3D rendered subject head & cortex. A slightly larger version can be found in (b) and an enlarged centered overview in (c) for all 100 MEPs, and thresholded (>0.1mV) version of the same data in (d). Each flag represents a stimulus nailed at the center of the TMS coil iso-field line intersecting with the cortex during each discharge. The rendering of the M1 area patch for ROI1 is shown in (e) and ROI2 in (f). Seed points for ROI creation are depicted as yellow spheres. Visualizations are obtained from SCIRun.

In the model validation described in the following sections, the performance of models using either ROI as a ‘MEP generating’ entity is compared, in the hope to observe a better fit for the true MEP generating region of the motor cortex. As MEPs can be evoked from the primary as well as the pre-motor cortex, we think it is important to test this assumption.

For patch isolation of ROI-1 and ROI-2, we developed in-house a new module to SCIRun (Modules::MiscField::SelectMeshROI), which filters triangles based on N-step topological distance from an initial seed element, depicted as a yellow sphere in figure 2.

#### 2.2.8 Activation Metrics

We introduced several different so-called ‘metrics’ in order to simulate the coupling between the injected current by TMS and neuronal activation in the cortical surface (and eventually recorded neuronal responses in the muscles by EMG). We looked into the total E-field as well as its tangential/orthogonal components in relation to the orientation of the local cortical sheet, using the two ROI patches described in the previous section. In order to construct a single scalar descriptive value from our multi-compartment volumetric model of induced currents, we always summed the resulting ‘activation’ over all polygons of the surface ROI divided by the total area spanned by it.

The activation ‘metric’ per surface area (which are triangles spanning the cortical surface here) is motivated by ideas of how assemblies of different neurons in the various layers of the cortical sheet are thought to be affected by external currents. Below we will explain and motivate all these metrics, which will be compared in their performance in explaining observed MEPs per stimulus.

The metric C0 was our zero hypothesis metric, which is electric field agnostic, and only depends on the distance from the coil to the target. The distance is simply the distance from centroids of triangles on ROI surface to the iso-center of the induced field of the coil (a line pointing downward from the figure of 8 coil center, perpendicular to the surface of the coil). Distance per surface triangle center is defined by the shortest distance from that point to the isocenter line of the coil.

The metric CE is motivated by the notion that many cells in the cortex respond to TMS-induced currents, and the mix of signals from the diverse population of neurons simply scales with the net magnitude of the induced E-field, irrespective of the angle with respect to the cortical sheet (perhaps because the interplay is too complex to model with such a crude metric).

Fox et al [17] introduced the popular C3 metric that is motivated by the columnar orientation of pyramidal cells in the cortical layers and assumes that an E-field perpendicular to the cortical surface evokes maximal action (in a symmetric manner, irrespective of whether the E-field points inward or outward from the surface). There is quite some indirect support for this metric [29], and also more recent criticism [6]. We introduced more metrics here reflecting the unidirectional current injections specific to our monophasic TMS device, namely C4 and C5 where both out- and in-wards currents are investigated independently. For completeness, we also introduced C2 which is simply the opposite of C3 proposition where we consider tangential injected current (parallel to the cortical surface) to be the most optimal in inducing neuronal activation. This could be the case when dis-inhibiting pyramidal cells through inter-neurons, generally extending parallel to the cortical surface, are an important driving factor in translating TMS induced E-fields to neuronal activation.

The metrics were computed as given in the formulas below, and are graphically depicted in figure 3. Our naive (null-hypothesis) metric, independent of the computed field was:

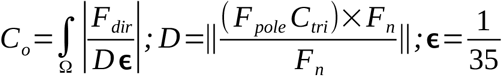 where Fdir is the direction of the flag (coil direction) (see figure 2), D denotes the shortest distance from a point to a line where Fpole is a point on the pole of the flag, *C_tri_* is the center of each triangle on the ROI patch, *F_n_* is the normalized ‘flag pole vector’ (the coil’s center line at the moment of TMS pulse delivery).

**Figure 3:**
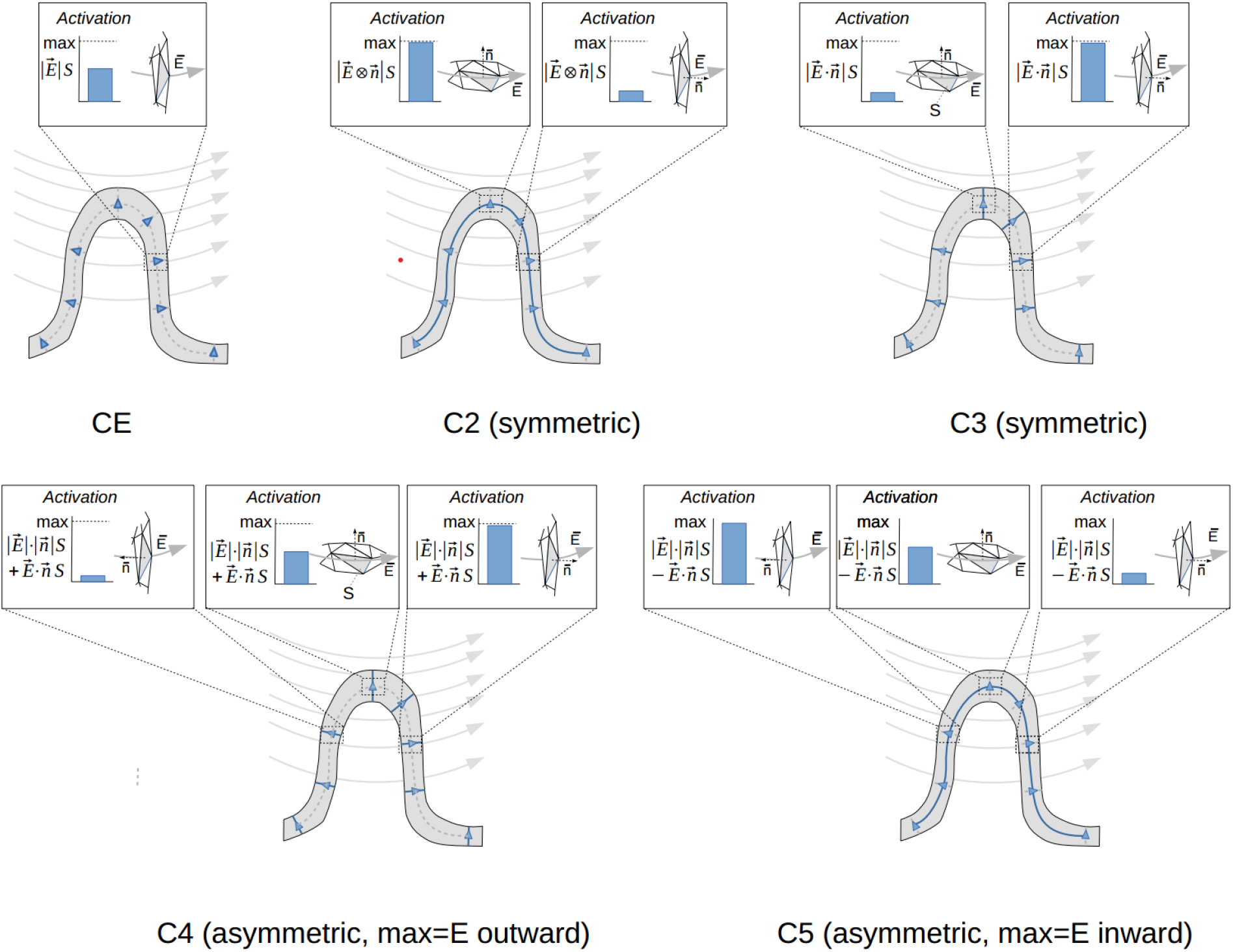
Conceptual illustration of the effect of each metric on the final estimate of the effectiveness of stimulation on a highly idealized folded cortical surface of a human cortex in 2D, for one gyrus in 3 situations. The curved grey lines denote the E-field E evoked by a TMS stimulus, S is the area of a surface patch, and n is the surface normals. The panels depict the geometrical relationship between a surface polygon normal n and the estimated E-field, given for the center of the polygon. **CE**: the activation per surface mesh element equals the size of the E-field times the area of the surface it passes through, independent from the current direction. **C3**: for each surface element, the resulting activation is the product of the surface area and the size of the vector product of the E-field and surface normal (size of both vectors times the cosine of the angle between them). This is maximal for a surface patch along the sulcal wall, and minimal for a surface patch on the gyral crown. **C2**: for each surface element, the resulting activation is the product of the surface area and the size of the cross product of the E-field and surface normal (size of both vectors times the sine of the angle between them). This is minimal for a surface patch along the sulcal wall, and maximal for a surface patch on the gyral crown. **C4** and **C5**: similar to C3 but flavoring outward or inward-injected E respectively.

The rest of the metrics CE,2,3,4,5 which take into account the magnitude of the E-field alone (CE) or in relation to the cortical surface orientation (C2-5), were formally defined as given below. Here **Et** is the total E-field (see section 2.2.4); **S** is the area of the polygon (triangle) and **n** is its normal vector (defining orientation). The preferential direction of the electric field for each of the metrics from C2 to C5 was also described after each formula.

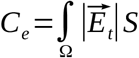 The E-field magnitude;
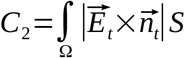 Symmetric E-field tangential;
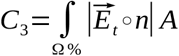 Symmetric E-field orthogonal;
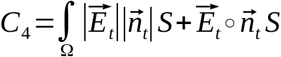 Asymmetric E-field orthogonal outward;
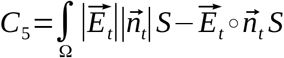 Asymmetric E-field orthogonal inward;

The integral over the domain is simply the sum over each triangle of the isolated tri-surface patches of ROI1 and ROI2, having each an area **S**. We considered the total electric field **Et** only. The ROI tri-surface patch geometries were exported to Matlab per ROI. The electric field was isolated for each patch in SCIRun and exported to Matlab. Then we computed each metric per patche and finally compared it against the quantified MEPs.

### 2.3 Experimental protocol

#### 2.3.1 Functional MRI of voluntary thumb movement

During fMRI scanning, the participant was instructed to perform thumb movements with the right thumb upon presentation of an auditory cue. The recording lasted 510 seconds. Thumb movements were measured with ECG electrodes integrated with the MRI system (effectively using them as EMG), attached to the thumb muscle.

#### 2.3.2 Concurrent TMS-EMG

For each subject, individual resting motor threshold (RMT) was determined, starting from 30% MO ramping up in steps of 5% until a detectable MEP was observed, both visually and via NeurosoftMEP software with the criteria of having more than 50mV peak-to-bottom MEP amplitude. Subsequently, intensity was reduced in steps of 1%, until we observed a successful MEP in 5 out of 10 trials [30]. Coil orientation was in AP direction while kept roughly orthogonal to the central motor gyri for each participant.

We conducted MEP mapping on predetermined subject-specific locations, organized roughly in a cross fashion. The procedure involved stimulation in a cross pattern on and around M1 (see figure 2). We relied on an image-based neuro-navigation system to achieve precise cortical targeting, where each coil placement was possible within 3mm spatial accuracy and with <5mm distance to target as a projection on the coil central iso-line (Brain Science Tools BV, The Netherlands). It involved 5 stimulation sites, one on top of M1 (which was determined from individual fMRI BOLD maximum, see section 2.3.1) as well as two locations about 1 cm more medial and lateral along the pre-central sulcus and two locations anterior and posterior from M1. See figure 2 for an impression. Each site was stimulated with 4 orthogonal coil orientations, each repeated 5 times, leading to a total of 100 stimuli (5 positions; 4 orientations; 5 repetitions). The TMS machine output was different for the stimulations directly on M1 (110%RMT) in contrast to the other 4 peripheral locations (120%RMT). This was done in an attempt to roughly match the intensity of the fields injected on all stimuli sites. This way the chance of supra-threshold minimum intensity would not bias in favor of the already anatomically predicted the best location. The orientation of the coil was also picked in a manner that the first orientation always pointed toward M1 ‘cross intersection’.

The order of stimuli was picked in a pseudo-random manner for each subject with a minimum of about 3 seconds inter-stimulus interval (to compensate for potential repetition suppression effects), around 10 seconds interval when changing between the 5 sites. The order of the four major coil orientations was picked in a similar manner per site. A typical session duration took around 15-20 minutes excluding preparation time.

Since our navigation system had an integration with the software package NeurosoftMEP of the EMG manufacturer, it is possible to have a comprehensive session export where each site of stimulation and raw MEP trace is recorded in a human-readable XML format. There are several important parameters exported per stimulus that are crucial for the correct placement of the coil model in the world space of our FEM simulations. The final rigid body transformation was derived as follows:, where (N – navigator matrix, based on a mapping between navigator and scanner world space; C – navigator coil generative space; S – navigator sensor tip offset; F1, F2 and are two flips to compensate for the difference in generative space between SCIRun and the navigation software where L coil thickness offset in Z+ direction along the coil iso-line). This was important to fully reconstruct each single coil position and orientation when running our FEM and activation metric model for each single stimulus (see section 2.4.3 for details), to obtain modeled MEP estimates.

### 2.4 Analysis

#### 2.4.1 Functional MRI and BOLD response

In order to obtain a subject-specific thumb area mapped on each participant’s motor cortex we had to post-process the anatomical T1-weighted MR images (see Section 2.2.7) and analyze the functional raw times MRI-EPI series of a trivial voluntary thumb movement task (see Section 2.3.1). For this purpose, we used the SPM12 software package [31], which is a freely available toolset for the commercially available Matlab 2014a MATLAB 2014a (The MathWorks, Inc., Natick, Massachusetts, USA). We considered the area corresponding to the thumb movement to be the location with the maximum BOLD response from the statistical activation map as given by SPM12. The statistical activation map was constructed based on event-related generalized linear model (GLM) analysis in a so-called event related analysis, where the thumb movement is modeled as a delta function convolved with the canonical hemodynamic response function (HRF). The timing of the thumb movements (and hence each delta function) was obtained from the EMG recordings that were acquired during MRI acquisition, using custom Matlab code. Two nuisance regressors were involved in the GLM analysis: the average BOLD signal in the WM and the CSF. The final statistical probability images were constructed based on a T-statistic with the T-threshold at P < 0.05, family-wise error (FWE) whole-brain corrected [31]. The maps of thumb movement activation around the hand knob in the pre-central gyrus were used to plan the stimulation sites around M1 (see section 2.2.7).

#### 2.4.2 Quantification of MEPs

We evaluated the recorded EMG traces for motor evoked potentials (MEPs) elicited by single-pulse TMS using a straightforward MATLAB routine assessing MEP amplitude as the peak-to-bottom signal difference in the interval 20ms after TMS pulse administration, where MEPs can be expected to occur. Our quantified MEPs were compared to the amplitudes computed by the NeuroMEP software package used to acquire the data, those measures are also based on the same peak to bottom criteria, and found to be nearly equivalent. One of the subjects exhibited relatively low MEP amplitudes (=< 0.5mV) and the automatic algorithm for onset-peak-bottom detection as part of the NeuroMEP software package failed to register any. This motivated us to write our own quantification routine in MATLAB as explained above, to have better control and ease of reproducibility of this analysis, rather than the MEP analysis provided by the NeuroMEP software.

#### 2.4.3 Descriptive Statistics

For each stimulus administered with TMS, the 6D position of the coil with respect to the head was stored in MRI native space coordinates by The Neural Navigator for later analysis. From this position, the incident field was computed using piece-wise Biot-Savart as explained above, and then FEM, the activation metrics over the surface patches around M1 were computed (which should be roughly proportional to MEP amplitude). This means we had a simulated and real MEP for each stimulus administered, which could be depicted against each other in a set of 2D points. We computed the correlation coefficient for each point cloud (that is, per metric and ROI) and subject.

Finally, the correlation coefficients were subjected to a 2-way repeated measures ANOVA, where we assessed whether one of the metrics and ROIs yielded a significantly different result compared to the others. If so, subsequent post-hoc tests were performed to test which of the combinations differed significantly. The significance level alpha was set to 0.05.

## 3 Results

### 3.1 Physiological EMG recordings

We first present the average observations for the MEPs recorded from the cross-shaped stimulation locations. In figure 4a, a set of sampled MEPs from one responsive position for one subject is presented. In figure 4-b, an overview of average MEP amplitudes for each location and direction is presented. The most optimal coil position seemed to be anterior of the motor hot spot (2.48mV) and the lateral of the motor hot spot (2.0mV) while the most optimal coil orientations were directed medially and anterior. The combination of M-M and A-A (medial location with medial direction and anterior location with anterior orientation, respectively) was most effective in evoking MEPs.

**Figure 4:**
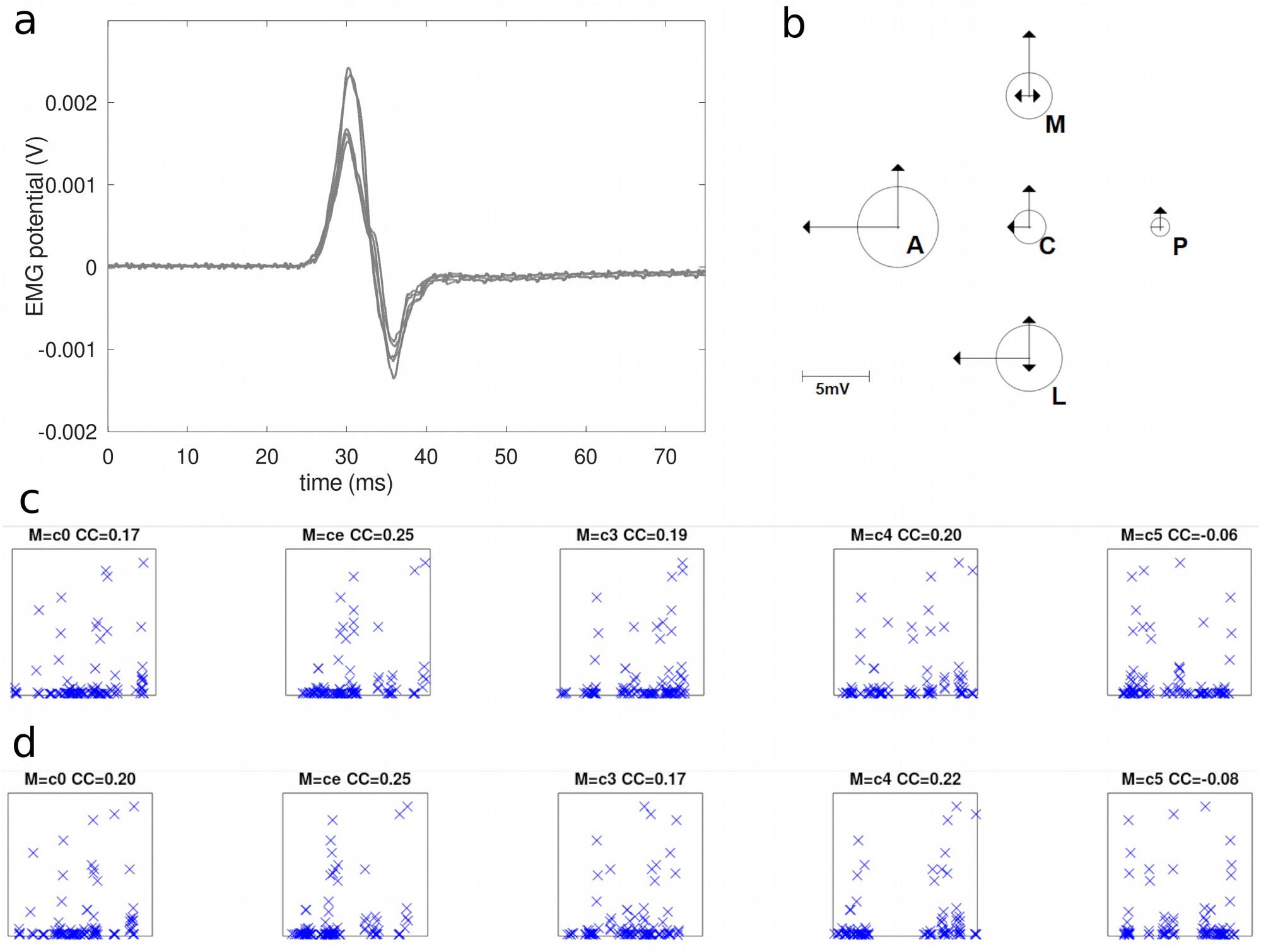
(a) Average MEP amplitudes are pooled over all subjects and shown separately for each position and orientation of the coil, in a schematic cross and directional arrow and circle graph. The radius of the circles shows the average MEP amplitude per stimulated location. The length of the arrows denotes the average MEP amplitude for each coil orientation for that location (note that for aesthetic reasons, arrowheads where not plotted when MEP amplitudes were less than 0.8mV). The stimulated location are detonated as A-anterior, P-posterior, M-medial, L-lateral, C-central according to convention and relative to the central hypothesized ideal hotspot determined from an individual fMRI activation. Orientations were relative to the central sulcus. (b) A subset of MEP traces for one representative subject (SU02), from location A with coil directed in the anterior direction, pointing away from the cenral sulcus. In this case, all stimuli repetitions caused a clear MEP response. (c) The scatter plots of modeled vs observed responses for subject (SU01) on ROI1. (d) The scatter plots of modeled vs observed responses for subject (SU01) on ROI2.

### 3.2 Simulations

We compared the quantified MEPs from the acquired EMG signals for each TMS stimulus, the peak to bottom magnitude, to the modeled ‘cMEP’. We calculated based on the suggested activation metrics, the sum/integral over the ROI surface patch weighted by the area (see section 2.2.8).

See figures 4-c and 4-d for the scatter plots of these cMEP x MEP pairwise comparisons, and the calculated correlation coefficients between those measures for one subject (SU06). Table 2 below presents the aforementioned correlation coefficients (CC) for all subjects, ROIs and activation metrics, and their mean and standard deviations. Metric C3 for ROI 1, and C5 for ROI2 had the highest correlation coefficient of modeled vs observed MEPs, i.e. the best correspondence (see table 2). This was confirmed by an omnibus ANOVA of ROI x metric (F(5, 45)= 8.51;p<0.005) implying the correlation coefficient was different for ROIs and metrics in general.

**Table 2:** Correlation coefficients for each metric evaluated per cortical ROIpatch on each subject individually.

| ROI<br>Metric | ROI 1 |  |  |  |  |  | ROI2 |  |  |  |  |  |
| --- | --- | --- | --- | --- | --- | --- | --- | --- | --- | --- | --- | --- |
|  | C0 | CE | C2 | C3 | C4 | C5 | C0 | CE | C2 | C3 | C4 | C5 |
| CC | 0,47 | 0,65 | 0,66 | 0,39 | 0,46 | -0,08 | 0,53 | 0,66 | 0,65 | 0,17 | 0,37 | 0,08 |
|  | 0,32 | 0,19 | -0,12 | 0,24 | 0,09 | 0,19 | 0,21 | -0,13 | -0,18 | 0,21 | -0,51 | 0,32 |
|  | 0,26 | 0,09 | 0,12 | 0,00 | 0,33 | -0,19 | 0,25 | 0,09 | 0,12 | 0,03 | 0,25 | -0,15 |
|  | 0,39 | 0,49 | 0,20 | 0,27 | 0,51 | -0,06 | 0,34 | 0,47 | 0,13 | 0,22 | 0,19 | 0,47 |
|  | 0,34 | 0,11 | -0,34 | 0,60 | 0,02 | 0,11 | 0,00 | -0,11 | -0,36 | 0,56 | -0,75 | 0,59 |
|  | 0,02 | -0,04 | -0,11 | 0,19 | -0,03 | -0,02 | 0,02 | 0,00 | -0,04 | 0,17 | -0,13 | 0,11 |
|  | -0,05 | 0,52 | -0,14 | 0,50 | 0,44 | 0,03 | 0,00 | 0,51 | 0,41 | 0,52 | 0,09 | 0,53 |
|  | 0,09 | 0,34 | 0,33 | 0,27 | -0,05 | 0,48 | -0,10 | -0,38 | -0,53 | 0,47 | -0,57 | 0,39 |
|  | 0,16 | 0,14 | 0,11 | 0,17 | 0,25 | 0,01 | 0,05 | 0,10 | 0,07 | 0,07 | -0,35 | 0,35 |
| Mean | **0,22** | **0,28** | **0,08** | **0,29** | **0,22** | **0,05** | **0,14** | **0,13** | **0,03** | **0,27** | **-0,16** | **0,30** |
| SD | 0,18 | 0,23 | 0,30 | 0,18 | 0,22 | 0,19 | 0,20 | 0,34 | 0,37 | 0,20 | 0,40 | 0,24 |

Post hoc paired sample one-sided T-tests were then performed testing (within each ROI) whether these C3 or C5 were indeed different from the C0 or CE, respectively (we tested one-sided as we expected directed metrics such as C3, C4, and C5 to outperform C0 and CE). C0 and CE are the more ‘banal’ zero-hypotheses that do not make any assumptions about the current direction or even current amplitude and are metrics we hoped to outperform. For ROI1, C3 was not significantly different from C0 or CE (T(8)=0.80;p=0.22 and T(8)=0.14;p=0.45, respectively). For ROI2, C5 was larger than C0 at trend level (T(8)=1.47;p=0.08) but not from CE (T(8)=1.17;p=0.13, respectively). In summary, when taken together, the metrics and ROIs yielded significantly different predictions, with C3 and C5 outperforming other metrics at first sight. However, when specifically tested against much more simplistic metrics C0 and CE, only C5 outperformed the simplest 0-hypothesis C0 at the trend level.

Although C0 was designed to be a naive ‘distance to target’ metric, it was not outperformed by the popular C3 metric, for neither ROI. For the larger ROI2, there was a trend of C5 outperforming C0, but not the other simple metric CE that only takes into account field magnitude at the neuronal interface.

There seems to be a tendency toward better model predictions for those metrics (C4 and C5) taking into account whether injected currents point inward or outward from the orientation of the local cortical sheet. At least, from the correlation coefficients reported in table 2, it is clear that CE and C3 are not forming better than the metrics C4 and C5 that take into account whether currents were directed inward or outward.

In figures 4-c and 4-d the simulated MEP is plotted against the measured MEP for 2 representative subjects. The same figures are provided for all participants in the supplementary material.

The correlation coefficient per metric, ROI and subject is presented in table 2 below, along with their means and standard deviations over subjects.

## 4 Discussion

We investigated finite element models of currents in individual heads, and ensuing neuronal activation patterns, induced by a typical TMS figure-of-8 coil by comparing model results with evoked EMG responses in the thumb of nine healthy volunteers. The lack of empirical validation in a growing field of macroscopic computational neuromodulation was one of the motivations to pursue this work.

We obtained reasonable explanatory power for some combinations of ROI and activation metric for about 3 individuals using mainly ROI1 and the C3 metric, and for 5 individuals for the ROI2 and the C5 metric. However, for the group of subjects as a whole, the results were not convincing. Taking into account more sophisticated measures such as coil details, individual head tissue characteristics, and several possible macroscopic neuronal activation metrics, did not systematically explain observed evoked responses better than relatively simple metrics such as CE, which simply reflects local total current magnitude. The main finding was that a FEM model and an activation metric that accounted for asymmetrically directed current interactions with the cortical surface (C5) performed best (at trend level) when accumulating over a larger ROI incorporating the motor cortex (ROI2). For this specific model, we obtained a moderate correlation between observed and modeled MEPs. The C5 metric over ROI2 performed almost equally well as the popular C3 metric over a smaller patch on the anterior bank of the central sulcus (ROI1), covering only primary motor cortex representations of the thumb. However, our rather uninformed metric C0, taking into account only the distance from the stimulated target to the ROI believed to result in MEPs, performed not much worse than C3, and was only outperformed by C5 over ROI2 at the trend level. The other rather naive metric CE taking into account current magnitude modeled with FEM but not the direction of the current relative to the cortical surface performed equally well to both C3 over ROI1 and C5 over ROI2. The other metrics performed less well. Overall, the metrics taking into account induced currents in some way (CE, C2, C3, C4, and C5) together performed better than the uninformed control metric C0, implying that computing currents, in general, are useful.

An important reason for the moderate level of success of combining TMS with EMG to validate results from computational models with observed MEPs is the notorious variability observed in subsequently evoked MEPs. To not make our experiments overly long and to not stress the participants too much, we used 5 repetitions per site where 20 now seems to be the recommended minimum[32].

In our scatter plots in figure 4 this is also clearly visible: while for some larger model-derived responses the individual measured responses tended to increase as well, there was a substantial number of recordings without responses at all, even for those stimuli where one would expect a large response. This might be a habituation effect of repeated TMS stimulation [12], or perhaps a modulation of motor responsiveness by other uncontrolled processes going on in the brain (state dependent) [33,34], which clearly reduced the effectiveness of our validation approach. The aforementioned papers on variability of MEPs largely appeared when the current experiments were already well underway, hence we could not take these recommendations into account. Nevertheless, simply increasing the number of repetitions might not be sufficient, as state-dependency of MEPs would continue to happen and reflect the inherent variability unrelated to low number of repetitions, and longer trains of identical stimulus might occasionally increase habituation effects.

Empirically, the strongest MEPs were not evoked from stimuli on the central site of the cross pattern of targets we used, focused on M1. Instead we observed strongest responses preferentially anterior to the primary motor cortex. In part this could be explained from the slightly lower intensity applied over the central crossing of our mapping pattern, which corresponds to the subject’s hotspot. Similar center-of-gravity (CoG) anterior displacements with respect to the anatomical primary motor cortex have been reported in various grid mapping studies before [9–11,35–37]. This has implications for the current practice of aiming the coil iso-center, which is assumed to be the focal point of stimulation for TMS, to a target of interest. We recently published a thorough study with a 5×5 grid mapping approach, observing the same phenomenon [38]. Another observation worth mentioning is the demonstrated capability of outward injected currents, that is TMS coil pointing away from the target of stimulation, to produce strong responses systematically. The combination of a Medial target and a Lateral to Medial current, and an Anterior and a posterior-anterior directed current were the two most efficient combinations of site and direction.

Our approach for empirically evaluating and optimizing TMS current models is by itself innovative and thorough, in our opinion. Unfortunately, the current data did not allow us yet to unequivocally select an optimal neuronal activation metric or cortical surface patch from which hand MEPs are supposed to be generated. Nevertheless, given the above, several improvements can be suggested for future work, to allow clearer distinction between the predictive value of detailed choices made in such models. Future studies should first and foremost adopt 20 to 30 repetitions per angulation and position of the coil to obtain an assessment of signal strength going into the corticospinal tract to produce an MEP, thus averaging out the uncontrolled variability that plagued our experiments. Even shorter sessions of successive stimuli, perhaps ten in a row, should be considered to avoid the notorious MEP habituation over repetitions, interleaved with brief stretching of hand muscles to avoid habituation. In the same light, a paradigm where location and angle are altered for every single stimulus, perhaps with a robot arm, might further reduce uncontrolled neuronal processes from becoming a dominant factor. This notion is supported by very recent work that demonstrated such randomizations can reduce the number of repetitions needed [39].

Finally, angle should be varied in steps of 45 degrees rather than 90 degrees as we used, to cover the probed domain in a bit more detail. Steps of 10 degrees are reportedly not changing results significantly [40] and should be considered the lower bound of angle step size.

With such optimizations of the experimental paradigm, it might be possible to make informed choices about how evoked currents induce neuronal activation, and from which part of the motor system, and perhaps also allow for other model optimizations [30,41].

## 5 Conclusion

Frameless stereotaxic neuronavigation for TMS in combination with computer-aided dosimetry has the potential to further improve the accuracy, reproducibility and general efficacy of clinical and experimental TMS. Such models would allow for optimization of the evoked current dose and activation pattern through adjusting key parameters, such as pulse intensity, coil placement, and orientation. However, our study challenges the notion that a simple metric based on the induced electric field and cortical surface orientation can predict TMS effects in the brain effectively given the current experimental approach.

There is no compelling reason at the moment to adopt simple metrics for neuronal activation as a result of our comparative analysis. The quality of the predicted motor evoked response was not much better than when using simpler assumptions such as total evoked current in any given region. For larger regions, there was some benefit of computing evoked currents over a mere distance-to-the-coil metric. The direction of the coil was not found to be a clear predictor of the resulting response either. Nevertheless, our experiments demonstrate that both were capable to deliver clear MEPs, and that for a subset of subjects we could explain a substantial part of the variance by taking into account the current direction.

Finally, future validation studies using EMG should adopt larger experiments with more repetitions of stimulation, and attempt to reduce MEP variability by adopting more optimal randomization of stimulus patterns. Current strategies, such as increasing repetitions, to achieve more robust recordings have the potential to miss subtle differences when it comes to the effects of coil orientation. This further contributes to making validation of FEM EM models of TMS-evoked activation using EMG even more challenging.

## Footnotes

1. SCIRun: A Scientific Computing Problem Solving Environment (Scientific Computing and Imaging Institute (SCI), University of Utah, USA). Additional modules that were used: coil geometry generator, BiotSavart thin-wire solver, ROI patch isolator; our own contributed module for coil models https://github.com/pip010/scirun4plus; FEM iterative solver based on the descending bi-conjugate gradient method with Jacobian per-conditioner.

2 BioMesh3D: Quality Mesh Generator for Biomedical Applications (Scientific Computing and Imaging Institute (SCI), University of Utah, USA). https://www.sci.utah.edu/devbuilds/scirun_docs/BioMesh3DGuide.pdf

3 Cleaver2: A Multi Material Tetrahedral Meshing Library and Application (Scientific Computing and Imaging Institute (SCI), University of Utah, USA). https://github.com/SCIInstitute/Cleaver2

